# Consumer-driven nutrient recycling of freshwater decapods: linking ecological theories and application in integrated multi-trophic aquaculture

**DOI:** 10.1101/2022.01.11.475807

**Authors:** Gabriela E. Musin, Maria Victoria Torres, Débora A. Carvalho

## Abstract

The Metabolic Theory of Ecology (MET) and the Ecological Stoichiometry Theory (EST) are central and complementary in the consumer-driven recycling conceptual basis. The comprehension of physiological processes of organisms at different levels of organizations is essential to explore and predict nutrient recycling behavior in different scenarios, and to design integrated productive systems that efficiently use the nutrient inputs through an adjusted mass balance. We fed with fish-feed three species of decapods from different families and with aquacultural potential to explore the animal-mediated nutrient dynamic and its applicability in productive systems. We tested whether physiological (body mass, body elemental content), ecological (diet), taxonomic and experimental (time of incubation) variables predicts N and P excretion rates and ratios across and within taxa. We also analysed body mass and body elemental content independently as predictors of N and P excretion of decapods across, among and within taxa. Finally, we verified if body content scales allometrically across and within taxa and if differed among taxa. Body mass and taxonomic identity predicted nutrient excretion rates both across and within taxa. When physiological variables were analysed independently, body size best predicted nutrient mineralization in both scales of analyses. Regarding body elemental content, only body P content scaled negatively with body mass across taxa. Results showed higher N-requirements and lower C:N of prawns than anomurans and crabs. The role of crustaceans as nutrient recyclers depends mainly on the species and body mass, and should be considered to select complementary species that efficiently use feed resources. Prawns need more protein in their feed and might be integrated with fish of higher N-requirements, while crabs and anomurans, with fish of lower N-requirements. Our study contributed to the background of MTE and EST through empirical data obtained from decapods and provided useful information to achieve more efficient aquaculture integration systems.

## Introduction

Consumers are important nutrient recyclers in aquatic ecosystems as a source or sink for elements such as carbon (C), nitrogen (N) and phosphorus (P) [1, 2]. The excretion and egestion of waste products are immediate processes by which animals can be a source of nutrients for primary producers and heterotrophic microorganisms [3]. Animals also constitute nutrient pools, as they grow and reproduce [1,2,4]. As stated by [5], “what animals eat and excrete shapes their role in ecological communities and determines their contribution to the flux of energy and materials in ecosystems”. In this sense, two ecological theories are central and complementary in the consumer-driven recycling conceptual basis [6]: the Metabolic Theory of Ecology (MTE) [7] and the Ecological Stoichiometry Theory (EST) [8]. Whereas one emphasizes on energy (MTE), the other does so on matter (EST) [5].

The MTE states that the rate at which organisms take up, transform, and expend energy and materials, is the most fundamental biological rate. Organisms are influenced by intrinsic (body size –hereafter, body mass– and stoichiometry of organisms) and extrinsic (temperature) factors, which, in turn, obey to chemical and physical principles [7]. According to the EST, the organismal response to food composition is a useful tool to predict the stoichiometric homeostasis of a given consumer through the selective retain and release (excretion and egestion) of elements like C, N and P. Rates and ratios by which animals recycle nutrients reflect the element imbalance between their bodies and their food [8]. The mass-specific excretion rate of nutrients (i.e. nutrients excreted per unit of body mass per unit time) generally decrease with increasing body mass due to allometric restrictions in metabolism [3,8–10]. A central concept in EST is the Growth Rate Hypothesis (GRH) [11, 8], which theorizes that differential allocation of body P content results from differential allocation of P-rich RNA required to protein synthesis during growth. Organisms with high rates of protein synthesis might have lower N:P content. The GRH also predicts that body P-content should be less in small-bodied organisms because these organisms tend to have faster growth rates.

[12] proposed a model that suggests that the N:P released by a homeostatic consumer increases with food N:P and decreases with body N:P content. Studies found mixed results in aquatic vertebrates and invertebrates, both supporting and contradicting this mass balance model [13–17]. The variable accuracy of diet and body content as predictors of N:P excretion has been attributed to the flexible homeostasis of organisms [15, 16], taxa-specific mechanisms [17], biotic and abiotic factors [18] and resource quality [19]. These mixed results must be due to taxa-specific metabolic processes. [20] found that body mass and taxonomic identity mainly predict excretion rates, but a better understanding of taxa-specific metabolic processes are necessary to predict animal-mediated nutrient recycling. Incubating animals in containers with a known volume of water is a common procedure to measure nutrient mineralization rates [13, 21]. However, time and conditions of incubation could also affect rate values and, therefore, experimental design should consider it [22].

Variation in body stoichiometry of organisms is hypothesized to be driven by taxon-specific constraints imposed by phylogeny [23–26], differential growth rates and allometry [8, 11], structural differences in material allocation [8,27,28], and trophic position [23,25,29]. In the last case, trophic position should predict body composition because organisms should minimize imbalances between the elemental body content and their diet. In this way, predators might have low C:nutrient and high N:P content because they have adapted better to ingest high-nutrient food than lower trophic-positioned organisms [24]. However, high C:nutrient could be found in organisms with C-rich structures, such as the chitin of the exoskeleton, due to differential nutrient allocation [8, 27]. In addition, variation in body composition might be due to ontogenetic changes in body stoichiometry and/or P demand associated with faster growth rates [28, 30]. Again, taxa-specific metabolic processes constraint the acquisition, incorporation, and release of elements by organisms [31] and, in doing so, scale of analysis may be important to perform any stoichiometric prediction. For example, [25] found family to be the best supported taxonomic level at which body content is conserved in a marine animal community, while [24], in detritus-based communities, and [26], in a global data-set, found that the level of taxonomy that best explains body content depends on the element in question.

The role of vertebrates [25,32,33] and invertebrates [31,34,35] as consumer-driven nutrient dynamics in aquatic systems has been extensively studied from different scales of analysis [20,23–26]. Among invertebrates of freshwater systems, decapod crustaceans were also target organisms, either as individuals, species or assemblies, or as agents that regulate the nutrient dynamics of freshwater ecosystems [17,18,28,36–40]. However, this crustacean order is still less evaluated than other taxa [2,28,38] despite the key function they fulfill in ecological process [41–44].

In lowland rivers in the Atlantic slope of southern South America, five families of decapods (Sergestidae, Palaemonidae, Parastacidae, Aeglidae and Trichodactylidae) belonging to four suborders (Dendrobranchiata, Caridea, Anomura and Brachyura) comprise the littoral-benthic communities of freshwater systems [45–48]. Despite the phylogenetic distance [49–52], differences in the ventrally folded pleon (carcinization) [53] and lifestyle [47], the trophic habit of these decapods is mostly omnivorous, generalist and opportunistic [46,48,54]. They fulfill key functions in the ecological processes of natural environments by crushing and processing decomposing plant material and by consuming aquatic invertebrates [42,44,54–56]. Although their trophic habits resemble the importance of vegetal and animal components, they could vary during ontogeny and interspecifically, in addition to extrinsic factors [57–59]. The trophic habit of these organisms is an advantage in captivity conditions because it allows a good acceptance of artificial feed and enables experimental aquaculture research [60, 61].

Nowadays, the accumulated information about these decapods is wide. They display different morphological and physiological traits among families, such as foregut types [55,62,63], predation strategies [64–66], enzyme and metabolic activity [61,67,68], and oxygen consumption [69], which certainly influence the resource acquisition, assimilation, and excretion. The holistic overview of this previous information, in addition to an experiment in a laboratory-controlled environment, could be a useful way to understand the taxa-specific differences in stoichiometry and metabolic rates (e.g. excretion) that influence the nutrient dynamic of natural and artificial aquatic systems.

In addition, there is an increasing interest in cultivating native species that are adapted to local conditions to conserve local biodiversity [70–72]. The use of native crustaceans in aquaculture systems is growing in South America [73–77]. The potentiality of using them as nutrient recycle organisms in fish production systems is high if we consider their wide trophic habits, the great acceptance of artificial feed, and the multiple uses of this by-product to add revenue to the production [60,78,79]. This approach is used to deal with the environmental burden of intensive farming and is coupled with the practice of integrated multitrophic aquaculture (IMTA), which consists in co-cultivating organisms of different trophic levels in the same system selected for their functions and complementary ecosystem services, connected by the transfer of matter and energy through water [80]. The low diversity of these artificial aquatic systems implies a strong influence of few taxa on the nutrient turnover [14, 81] and emphasizes the importance of studying species with aquacultural potential and interest.

Coupling experimental aquaculture research with ecological theories is an interesting way to explore the animal-mediated nutrient dynamic and its applicability in productive systems. Here we test how physiological (body mass, body elemental content), ecological (diet), taxonomic and experimental variables explain nutrient recycling at order (across taxa) and at family/species (among and within taxa) levels, in order to discuss them from an ecological and applied point of view. We used three species of decapods from different suborders and families, with antecedents about trophic and digestive ecology and with aquacultural potential use. We are interested in understanding their potential nutrient recycle role in natural and artificial systems, such as in an IMTA. We performed an experimental research using two different commercial feedstuffs elaborated for detritivorous and omnivorous fish, and we analysed the excretion response variables (mass-specific excretion rates of N, P and associated ratios) in two different incubation’s period. We also evaluated the body elemental content of C, N, P and associated ratios to relate these results with antecedents about trophic habits, carcinization and ontogeny.

We asked three main questions **(Q)**. **Q1:** Which of the physiological (body mass, body elemental content), ecological (diet), taxonomic and experimental (time of incubation) variables best predicts N and P excretion rates and ratios across and within taxa? We hypothesized that: **H1-** Physiological and taxonomic variables best explain the variation in N and P excretion rates. **Q2: a)** Could physiological variables (body mass, body elemental content), if analysed independently, predict N and P excretion rates and ratio of decapods, according to MTE and EST? Are these allometric (excretion rate vs. body mass) and stoichiometric (excretion rate vs. body elemental content) variations observed across and within taxa? **b)** Are allometric and stoichiometric variations different among taxa? With respect to these questions, we formulated three hypotheses: **H2-** N:P excretion rates decrease with increasing N:P body content, as postulated by Sterner (1990), both across and within taxa. **H3-** N and P excretion rates scale allometrically across and within taxa. **H4-** There are allometric and stoichiometric differences among taxa. **Q3: a)** Do body content scales allometrically across and within taxa? **b)** Are body elemental content and allometric variations different among taxa? We hypothesized that: **H5-** Body P content is negatively related to body mass across and within taxa, according to GRH. **H6-** Body C:N content increases in carcinized crustaceans and decreases, together with N:P, in lower trophic positioned species.

## Material and methods

### Studied organisms

Decapod species used in this experiment represent three different families of neotropical decapod crustaceans. *Macrobrachium borellii* is a prawn of the Palaemonidae family, with wide distribution in La Plata Basin of northern Argentina, Paraguay and southern Brazil [82, 83]. Its natural diet exhibits a significant presence of animal items such as dipterans and oligochaeta larvae, and a low importance of vegetal remains and algae [47, 54, 56]. It is a species characterized by moving in the water column and towards the littoral vegetation [62, 84]. *Aegla uruguayana* belongs to the Aeglidae family, which has a unique genus endemic to southern South America. This species presents benthic habits, typically sheltered at the bottom and under rocks of current rivers and streams [82, 85] and displays strong swimming habits [86, 87]. Its natural diet is composed mainly of vegetal remains and diatoms with low importance of animal items [54, 55]. *Trichodactylus borellianus* belongs to the neotropical family of freshwater crabs, Trichodactylidae, with broad distribution in South America (from 0° to 35° S) [88]. This crab has a close relationship with the floating aquatic vegetation and exhibits little mobility [48, 89]. Its natural diet is characterized by vegetal remains, algae and animal items such as oligochaetes and insect larvae [90, 56]. Both prawns and crabs form abundant populations associated with the aquatic macrophytes of the floodplain littoral zone [46,89,91].

### Crustaceans sampling and laboratory maintenance

Specimens of crustaceans of varied sizes were manually collected from the environment with the aid of a hand net (500 µm mesh size) during the austral late spring (November and December 2018). *Macrobrachium borellii* and *T. borellianus* were captured from the “Ubajay” stream (31°33’43.45’’S, 60°30’58.73’’W), Santa Fe (Argentina), among the aquatic vegetation at the shoreline of water bodies. *Aegla uruguayana* were captured from “El Espinillo” stream (31°47’09.16’’S, 60°18’57.46’’W), Entre Ríos (Argentina), through the removal of stones at the bottom of the stream and placing the hand net against the current. Crustaceans were translated to laboratory in plastic containers, where they were acclimated gradually (at least two weeks) to the experimental conditions (temperature - 24 ± 1 °C; natural photoperiod – dawn and dusk around 05:00 and 20:00, respectively; conductivity– 250 ± 10 µS) in aquaria with dechlorinated and aerated tap water, with rocks and PVC tubes as refugees. During this period, crustaceans were fed *ad libitum* with the same fish feed used in the experiments. Two extruded commercial feeds (Garay SRL) were used. They were elaborated for the nutrition of omnivorous (OF) and detritivorous fish (DF) (e.g. pacú - *Piaractus mesopotamicus* and sábalo - *Prochilodus lineatus*) (Table 1). Both feeds were ground in a mortar and passed through sieves of 1000-µm-diameter mesh to attain a size that facilitates the ingestion by crustaceans. Then, feeds were stored in glass bottles at 5°C. Every 48 hours, feces and food remains were removed by siphoning and the discharged water was supplemented. Every day pH, conductivity, dissolved solids and temperature were measured with a waterproof tester (Hanna HI98129, Romania), and dissolved oxygen with an oximeter to verify water quality (YSI Proodo SKU626281, USA).

**Table 1.** Proximal and elemental (C, N, P) composition of detritivorous (DF) and omnivorous (OF) fish feeds used in the experiments.

| Ingredients | OF | DF |
| --- | --- | --- |
|  | (%) | (%) |
| Proximal composition (g/100 g dry basis)<sup>1</sup> |  |  |
| Protein | 27 | 30.68 |
| Lipids | 3.87 | 5.00 |
| Humidity | --- | 3.82 |
| Crude Fiber | 3.53 | 12.24 |
| Ash | 4.28 | 8.37 |
| P total | 0.80 | 1.24 |
| Elemental composition (g/100 g dry basis) |  |  |
| C | 42.4±0.3 | 41.4±0.1 |
| N | 3.8±0.1 | 4.9±0.2 |
| P | 1.1±0.2 | 1.6±0.2 |

<sup>1</sup> Garay SRL (Recreo, Argentina) – nutritional values analyzed by the manufacturer. --- not informed by the manufacturer

### Experimental design and procedures

The feeding experiment was carried out using specimens of each species with variable body mass and control for each treatment. Only *A. uruguayana* had a reduced number of specimens in the experiment due to the limited wild caught species. Feeds used were commercial formulations designed for the nutrition of fishes with omnivorous and detritivorous feeding habits (Garay SRL, Argentina).

After the acclimation period, organisms were transferred to individual plastic recipients of 1 liter, randomly and interspecified arranged, where they were left for 24 hours without food. In total, 66 individuals were used (24 of *M. borellii*, 18 of *A. uruguayana* and 24 of *T. borellianus*) plus six controls without specimens. Each recipient was provided with shelter (a small rock previously washed and chlorinated), artificial aeration and covered with a half shadow to reduce stress and prevent escapes. After the fast period, fish feed was offered *ad libitum* to the corresponding treatment and left for 90 minutes. Then, each organism was removed from the plastic recipients, washed with distilled water, and transferred to glass bottles with 150 ml of filtered (MG-F 0.7 µm, Munktell Filter-Sweden) and dechlorinated tap water. Then, 15 ml of water samples were taken from each recipient at 30 and 60 minutes using a micropipette (5000 µl), and they were conserved at −20°C until the analytical determination of excreted nutrients was performed. The incubation time of 60 minutes was previously considered adequate because excretion rates rapidly decrease after organisms stop the feed ingestion [92]. However, water samples were taken in each recipient at 30 and 60 minutes to verify if these incubation time lapses (also mentioned as time) represent a variation in nutrient excretion rates and associated ratios.

At the end of the experiment, crustaceans were kept without feed for 24 hours to eliminate the gut content. Then, animals were anesthetized in cold water and frozen. Subsequently, individuals were oven dried at 50 °C to constant weight, and weighted (± 1 µg) to determine dry body mass. The dried body of each individual was pulverized and homogenized in a mortar, and elementally analysed (C, N and P). This study adheres to the ethical standards of [93], and ethical and legal approval was obtained prior to the start of the study by the committee of ethics and safety in experimental work of CONICET (CCT, Santa Fe).

### Analytical and elemental analysis

Water samples from each treatment were analysed for inorganic forms of N and P, ammonium (NH_4_-N) and orthophosphate (P-PO_4_), respectively. NH_4_-N was quantified through the indophenol blue method [94], and P-PO_4_ through the ascorbic acid method [95]. To determine the stoichiometric proportion of carbon (C) and nitrogen (N) of crustaceans and feeds, two subsamples of each sample were elementally analysed in a CHN628 Series Elemental Determinators (LECO ®). For total phosphorus (P) analysis, dry crustaceans’ bodies or ground powder feed were weighted (± 1 µg) and combusted in a muffle furnace at 550°C for a minimum of 2 hours. Then, the mass of ash was weighted and acid-digested with 25 ml of HCl 1N during 15-20 minutes in a heating plate. The digested solution was brought to 100 ml with distilled water [96] and analysed using the ascorbic-acid method [95]. C, N and P of feed and crustaceans’ bodies were expressed as g/100g at a dry weight basis (dw) and ratios C:N, C:P and N:P were calculated using molar values.

The amount of ammonium and orthophosphate obtained in each excretion chamber was corrected by subtracting the average of nutrient concentration obtained in the control replicates (chambers without crustaceans) from the value obtained in crustaceans’ excretion chambers after 30 and 60 minutes. Then, results were divided by the body mass (mg of dry weight) and by the time of incubation (0.5 or 1 hour). Mass-specific NH_4_-N, P-PO_4_ and NH_4_-N:P-PO_4_ excretion rates were expressed throughout the text as N, P and N:P excretion or mineralization.

### Data analysis

The differences of elemental composition between two extruded commercial feeds for OF and DF were analyzed through Mann-Whitney-Wilcoxon Test with R software version 3.6.3 [97].

#### Nutrient excretion rates

**Q1-** Mixed-effects models with individuals as a random variable (lmerTest R package) [98] were used to analyse the effect of variables on the N, P and N:P excretion across and within taxa. The ‘Drop1’ function was run on the mixed-effects models to select the variables that best explain nutrient excretion rates and ratios variations. This function computes likelihood ratio test statistics and p-values for all single terms, fits those models and computes a table of the changes in fit. Only relevant interactions then compose the statistical table. The ‘single term deletion’ is useful for model selection and tests of marginal terms [98].

Mixed models were chosen, starting with the interaction effects of species variable (also named as taxonomic identity) (only for across taxa analysis), time and type of feed as factors, and body mass and body elemental content as covariates. Individuals were included as random effects to assess how time affects nutrient rates. If the term time was statistically significant, the data with higher nutrient excretion rates will be selected to run the posterior regressions and analysis of covariance (ANCOVA).

**Q2-a)** After running mixed models, allometric relationships across and within taxa were analysed between N, P and N:P excretion for 30 or 60 minutes by regressing these parameters as a function of species body mass. Also, stoichiometric relationships were quantified, across and within taxa, regressing nutrient excretion rates and ratios as a function of body N, P and N:P content. **b)** Allometric and stoichiometric trends among taxa were compared using ANCOVA analysis, including nutrient excretion rates and ratios (data of 30 or 60 minutes according to lmer results) as dependent variables with body mass and body elemental content (N, P and N:P) as covariates, respectively. Statistical differences among taxa were determined using Bonferroni adjustment for multiple testing corrections with the function emmeans_test of emmeans package [99] (Lenth et al., 2021).

#### Body elemental content

**Q3-a)** Allometric relationships across and within taxa were analysed between body N, P, C content and associated ratios by regressing these parameters as a function of species body mass. **b)** The body content of nutrients and ratios were compared among taxa through ANCOVA and using body mass as covariate. Statistical differences among species were determined using Bonferroni adjustment for multiple testing corrections with the function emmeans_test of emmeans package [99].

All analyses of nutrient excretion rates and body elemental content were conducted with R software version 3.6.3 [97] and all variables used were log_10_-transformed to standardize variance among species.

Finally, the relationship between log_10_-transformed physiological variables (body mass and body elemental content) and nutrient excretion was shown through Principal Component Analysis (PCA), based on a correlation matrix, as a conclusion and visualization of the main results. This analysis included nutrient excretion rates and ratios of the selected time (30 or 60 minutes), body elemental contents (single elements and ratios), and body mass, considering the taxonomic identity of each species.

## Results

The elemental composition analysis determined that DF presented a slightly higher amount of nitrogen (4.9 ± 0.2%) and phosphorus (1.6 ± 0.2%) compared to the OF (3.8 ± 0.1% and 1.1 ± 0.2%, respectively), but these differences were not significant (W= 0, *p=* 0.2 and *p=* 0.1, respectively). Regarding the percentage amount of carbon, both feeds were similar (W: 6; *p*= 0.2) (Table 1). Mean body masses of decapods used in the experiments were: *M. borellii* (351.0 ± 82.1 mg dw), *A. uruguayana* (554.2 ± 536.8 mg dw) and *T. borellianus* (110.8 ± 50.7 mg dw), with ranges of [44.3 - 377.6 mg dw], [8.2 – 1900.1 mg dw] and [26.7 – 220.9 mg dw], respectively.

### Nutrient excretion

**Q1-** Across taxa analysis identified species (taxonomic identity) and body mass as the variables that best explained N and P excretion, whereas only taxonomic identity explained variations in N:P excretion (Table 2). The type of diet and body N, P, and N:P content did not affect variations of nutrient excretion rates and ratio (Table 2). Moreover, time and its interaction with the taxonomic identity influenced the variation of N excretion (Table 2). Excretion levels of N were higher at 30 minutes (0.9574 ± 1.1097 µg N mg dw^-1^.hr^-1^) than at 60 minutes (0.7279 ± 0.7373 µg N mg dw^-1^.hr^-1^). According to these results, posterior regressions and ANCOVAs analyses were made using the data obtained at 30 minutes.

**Table 2.** Results of the variables selected by ‘Drop1’ function to explain the nutrients excretion rates and ratios, ran on the mixed-effects model across taxa.

|  | F-value | p-value |
| --- | --- | --- |
| N excretion |  |  |
| Time: Species | 5.4434 | 0.0096 ** |
| Time: Diet | 1.1026 | 0.3020 |
| Species: Diet | 1.2891 | 0.2925 |
| Time | 8.2662 | 0.005643 ** |
| Species | 42.985 | 5.241x10<sup>-12</sup> *** |
| Diet | 1.4157 | 0.2392 |
| Body mass | 19.7381 | 0.0001 *** |
| Body N content | 0.3992 | 0.5330 |
| P excretion |  |  |
| Time: Species | 1.1041 | 0.3408 |
| Time: Diet | 0.6410 | 0.4282 |
| Species: Diet | 0.4949 | 0.6129 |
| Time | 1.6150 | 0.2096 |
| Species | 17.7731 | 1.006x10<sup>-06</sup> *** |
| Body mass | 5.4282 | 0.0237 * |
| Body P content | 0.1090 | 0.7430 |
| N:P excretion |  |  |
| Time: Species | 0.6922 | 0.5108 |
| Time: Diet | 0.6355 | 0.4336 |
| Species: Diet | 1.8923 | 0.1608 |
| Time | 0.1359 | 0.7139 |
| Species | 39.0418 | 2.226x10<sup>-11</sup> *** |
| Body mass | 0.0236 | 0.8793 |
| Body N:P content | 0.0814 | 0.7781 |

Statistically significant differences \* $p < 0.05$ , \*\* $p < 0.005$ , \*\*\* $p < 0.001$ .

According to taxa analysis, the variables that most affect N excretion of the prawn *M. borellii* were body mass and body N content (Table 3), while any variable explained P excretion variations in this species (Table 3). Body mass explained the N and P excretion variations of *A. uruguayana* (Table 3). Moreover, time influenced the N excretion of this species (Table 3), with values higher at 30 minutes (0.9553 ± 1.6341 µg N mg d w^-1^.hr^-1^) than at 60 minutes (0.3128 ± 0.3542 µg N mg d w^-1^.hr^-1^). *T. borellianus* crab did not show a significant relationship between N and P excretion with any analysed variables (Table 3). Additionally, any variable explained N:P excretion variations in any species (Table 3).

**Table 3.** Results of the variables selected by ‘Drop1’ function to explain the nutrients excretion rates and ratios, run on the mixed-effects model within taxa.

|  | F-value | p-value |
| --- | --- | --- |
| Excretion N |  |  |
| *M. borellii* |  |  |
| Time: Diet | 1.3056 | 0.2798 |
| Time | 0.3264 | 0.5792 |
| Diet | 0.0502 | 0.8282 |
| Body mass | 8.9670 | 0.0172 * |
| Body N content | 6.2851 | 0.0365 * |
| *A. uruguayana* |  |  |
| Time: Diet | 3.2176 | 0.1030 |
| Time | 19.7189 | $3 \times 10^{-4}$ *** |
| Diet | 1.3291 | 0.2671 |
| Body mass | 9.1818 | 0.0163 * |
| Body N content | 0.1982 | 0.6679 |
| *T. borellianus* |  |  |
| Time: Diet | 0.0486 | 0.8311 |
| Time | 0.1212 | 0.7320 |
| Diet | 1.4188 | 0.2521 |
| Body mass | 1.6631 | 0.2447 |
| Body N content | 1.3292 | 0.2928 |
| Excretion P |  |  |
| *M. borellii* |  |  |
| Time: Diet | 0.3710 | 0.5504 |
| Time | 4.0167 | 0.0596 |
| Diet | 1.1349 | 0.2999 |
| Body mass | 0.0087 | 0.9267 |
| Body P content | 0.0596 | 0.8101 |
| *A. uruguayana* |  |  |
| Time: Diet | 2.1144 | 0.1715 |
| Time | 1.8092 | 0.1974 |
| Diet | 0.7377 | 0.4031 |
| Body mass | 15.3083 | 0.0020 ** |
| Body P content | 0.0175 | 0.8969 |
| *T. borellianus* |  |  |
| Time: Diet | 2.5337 | 0.1366 |
| Time | 0.0000 | 0.9947 |
| Diet | 1.4253 | 0.2401 |
| Body mass | 0.7611 | 0.4017 |
| Body P content | 0.2193 | 0.6487 |
| Excretion N:P |  |  |
| *M. borellii* |  |  |
| Time: Diet | 1.5418 | 0.2425 |
| Time | 0.4891 | 0.4938 |
| Diet | 0.0328 | 0.8585 |
| Body mass | 0.6743 | 0.4351 |
| Body N:P content | 0.1420 | 0.7168 |
| *A. uruguayana* |  |  |
| Time: Diet | 2.7509 | 0.1544 |
| Time | 0.3512 | 0.5604 |
| Diet | 1.5098 | 0.2342 |
| Body mass | 0.0097 | 0.9254 |
| Body N:P content | 0.9287 | 0.3637 |
| *T. borellianus* |  |  |
| Time: Diet | 0.4942 | 0.5129 |
| Time | 0.1916 | 0.6768 |
| Diet | 4.1843 | 0.1165 |
| Body mass | 0.5910 | 0.4862 |
| Body N:P content | 0.1285 | 0.7400 |

Statistically significant differences \* $p < 0.05$ , \*\* $p < 0.005$ , \*\*\* $p < 0.001$ .

**Q2-a)** According to regressions performed at 30 minutes and across taxa, N and P excretion rates decreased significantly with increasing decapods body mass. Decapods’ body mass explained more variation in N (45% of variance) than in P excretion (13% of variance) (Fig 1a, b). N:P excretion did not show significant differences with crustaceans’ body mass (Fig 1c). The stoichiometric relationship between N excretion and body N content was significant and positive, with an explanation of 36% (Fig 1d), whereas P excretion did not show significant differences with body P content (Fig 1e). N:P excretion decreased significantly with increasing body N:P content, with a low explanation (18% of variance) (Fig 1f).

**Fig 1.**
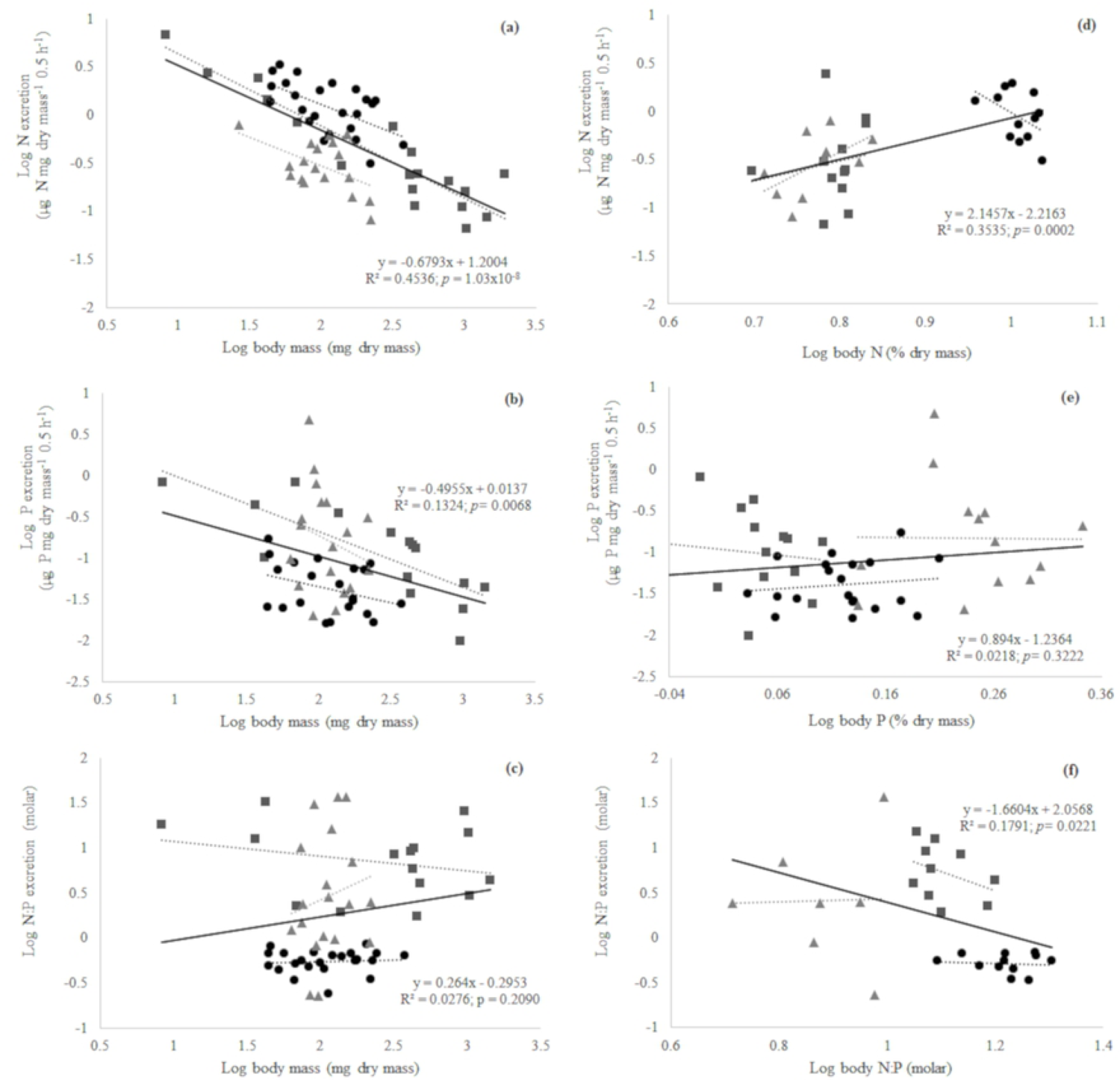
Linear regressions across and within taxa of excretion rates of N, P and N:P as a function of invertebrate body mass (a, b, c) and body elemental content (d, e, f). The equation, R-squared and p-value belong to the across taxa linear regressions. *Macrobrachium borellii* (black circle), *Aegla uruguayana* (dark gray square), *Trichodactylus borellianus* (light gray triangle).

Regarding regressions carried out at 30 minutes within taxa, N excretion of prawn *M. borellii* decreased significantly with increasing prawn body mass although it showed a low allometric relationship (Table 4) (Fig 1a). This prawn did not show a significantly linear relationship with P excretion and body mass (Table 4) (Fig 1b). *A. uruguayana* aeglid showed a significant and negative allometric relationship between N and P excretion and body mass, and this relationship was stronger in N than in P excretion (Table 4) (Fig 1a, b). *T. borellianus* crab showed a significant linear relationship between N excretion and body mass, but this was on the edge of significance (Table 4) (Fig 1a). This crab showed no significant linear relationship between P excretion and body mass (Table 4) (Fig 1b). According to N:P excretion, this variable did not show significant linear relationship with body mass in any species of decapods (Table 4) (Fig 1c). On the other hand, there was no significant linear stoichiometric relationship between nutrient excretion and body elemental content in any species of decapods (Table 4) (Fig 1d, e, f).

**Table 4.** Within taxa linear regressions of nutrient excretion rates (N, P) and ratio (N:P) as a function of crustaceans’ body mass.

|  | Slope (b) | R<sup>2</sup> | p-value |
| --- | --- | --- | --- |
| N excretion vs body mass |  |  |  |
| *M. borellii* | **-0.5887** | **0.3430** | 0.0026 ** |
| *A. uruguayana* | **-0.7518** | **0.8544** | 4.25x10<sup>-08</sup> *** |
| *T. borellianus* | **-0.5628** | **0.2203** | 0.0426 * |
| P excretion vs body mass |  |  |  |
| *M. borellii* | -0.3978 | 0.1288 | 0.1100 |
| *A. uruguayana* | **-0.6783** | **0.6179** | 5x10<sup>-4</sup>*** |
| *T. borellianus* | -0.9372 | 0.0547 | 0.3570 |
| N:P excretion vs body mass |  |  |  |
| *M. borellii* | 0.0395 | 0.0073 | 0.6920 |
| *A. uruguayana* | -0.1600 | 0.0656 | 0.3568 |
| *T. borellianus* | 0.7526 | 0.0299 | 0.5340 |
| N excretion vs body N |  |  |  |
| *M. borellii* | -5.1430 | 0.2069 | 0.1374 |
| *A. uruguayana* | 2.0270 | 0.0261 | 0.6155 |
| *T. borellianus* | 4.5670 | 0.3190 | 0.0889 |
| P excretion vs body P |  |  |  |
| *M. borellii* | -3.0430 | 0.0497 | 0.5639 |
| *A. uruguayana* | -1.4063 | 0.0116 | 0.7129 |
| *T. borellianus* | -2.1786 | 0.0112 | 0.6954 |
| N:P excretion vs body N:P |  |  |  |
| *M. borellii* | -0.1773 | 0.0107 | 0.7485 |
| *A. uruguayana* | -2.1400 | 0.1334 | 0.2995 |
| *T. borellianus* | 0.1442 | 0.0004 | 0.9640 |

Statistically significant linear relationships between species body mass and the respective response variable \* $p < 0.05$ , \*\* $p < 0.005$ , \*\*\* $p < 0.001$ .

**b)** Among taxa, it was observed a significant individual effect of taxonomic identity and body mass (and not interactive effect) on N (ANCOVA, *p=* 2.79 x 10^-12^ for species effects and *p=* 2.84 x 10^-14^ for body mass effect) and P (ANCOVA, *p=* 0.0004 for species effects and *p=* 0.0002 for body mass effect) excretion. N:P excretion showed a significant difference only between taxonomic identity and not as regards body mass (ANCOVA, *p=* 8.29 x 10^-10^ and *p=* 0.640).

When body elemental content was used as a covariate, it was observed a significant individual effect of taxonomic identity and body N content (and not an interactive effect) on N excretion (ANCOVA, *p=* 0.0014 and *p=* 0.0003, respectively). Whereas in P and N:P excretion, there was only a significant individual effect of taxonomic identity (ANCOVA, *p=* 0.0094 and *p=* 1.03 x 10^-6^) and not of body elemental content.

Regarding *post hoc* tests, N excretion was statistically different among all species (Bonferroni *post hoc*, *p*< 0.001) (Fig 2). *M. borellii* prawn showed the highest average value of N excretion, *A. uruguayana* showed an intermediate one, and *T. borellianus* the lowest one (Fig 2a). P excretion was statistically different among *M. borellii* prawn and the other species (Bonferroni *post hoc*, *p*< 0.001) (Fig 2b). This species showed lower average value of P excretion than the other species (Fig 2b). The N:P excretion differed significantly among all species (Bonferroni *post hoc*, *p*< 0.001) (Fig 2c). *M. borellii* prawn showed the lowest average value of N:P excretion, *A. uruguayana* showed the highest one, and *T. borellianus* an intermediate one (Fig 2c).

**Fig 2.**
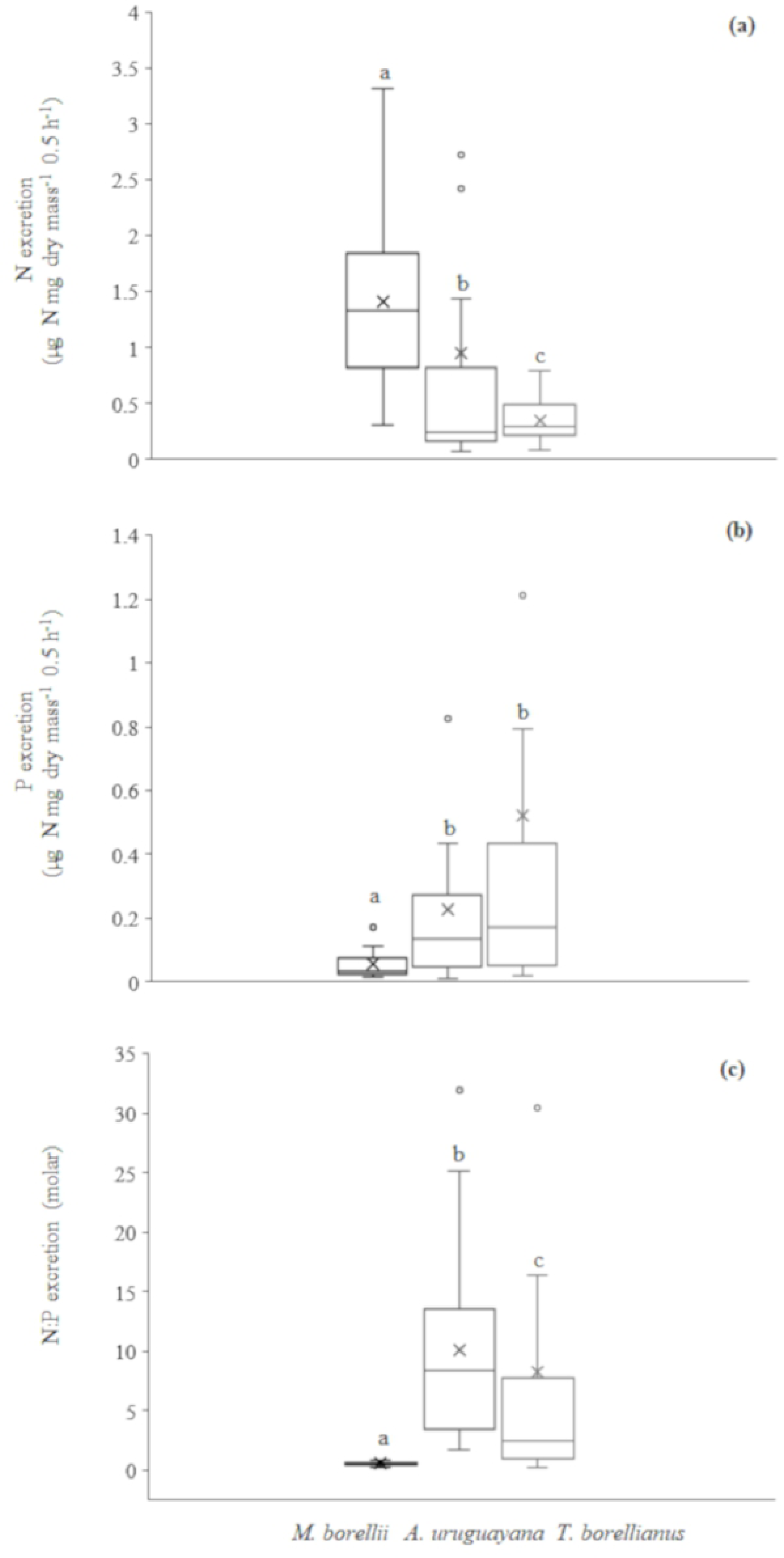
Box plots of excretion rates of N, P and N:P of each decapod species (a, b, c). The top, bottom, and line through the middle of the boxes correspond to the 75th, 25th, and 50th (median) percentile. The whiskers extended from the 10th percentile to the 90th percentile. Crosses indicate the median values. Different letters above bars indicate significant differences among taxa (*p* < 0.05).

### Body elemental content

**Q3-a)** Regressions across taxa showed a significant negative allometric relation between body P content and body mass (Fig 3b). Whereas the other body content elements and ratios did not show a significant relation with body mass (Fig 3a, c, d, e, f). As regards taxa regressions, only *T. borellianus* crab showed a significant negative allometric relationship between body N content and body mass (Table 5) (Fig 3a). Variations in body P and C content did not show significant linear relationship with body mass in any species of decapods (Table 5) (Fig 3b, c). Only *M. borellii* prawn and *T. borellianus* crab showed a significant positive linear relationship between body C:N content and body mass (Table 5) (Fig 3e).

**Fig 3.**
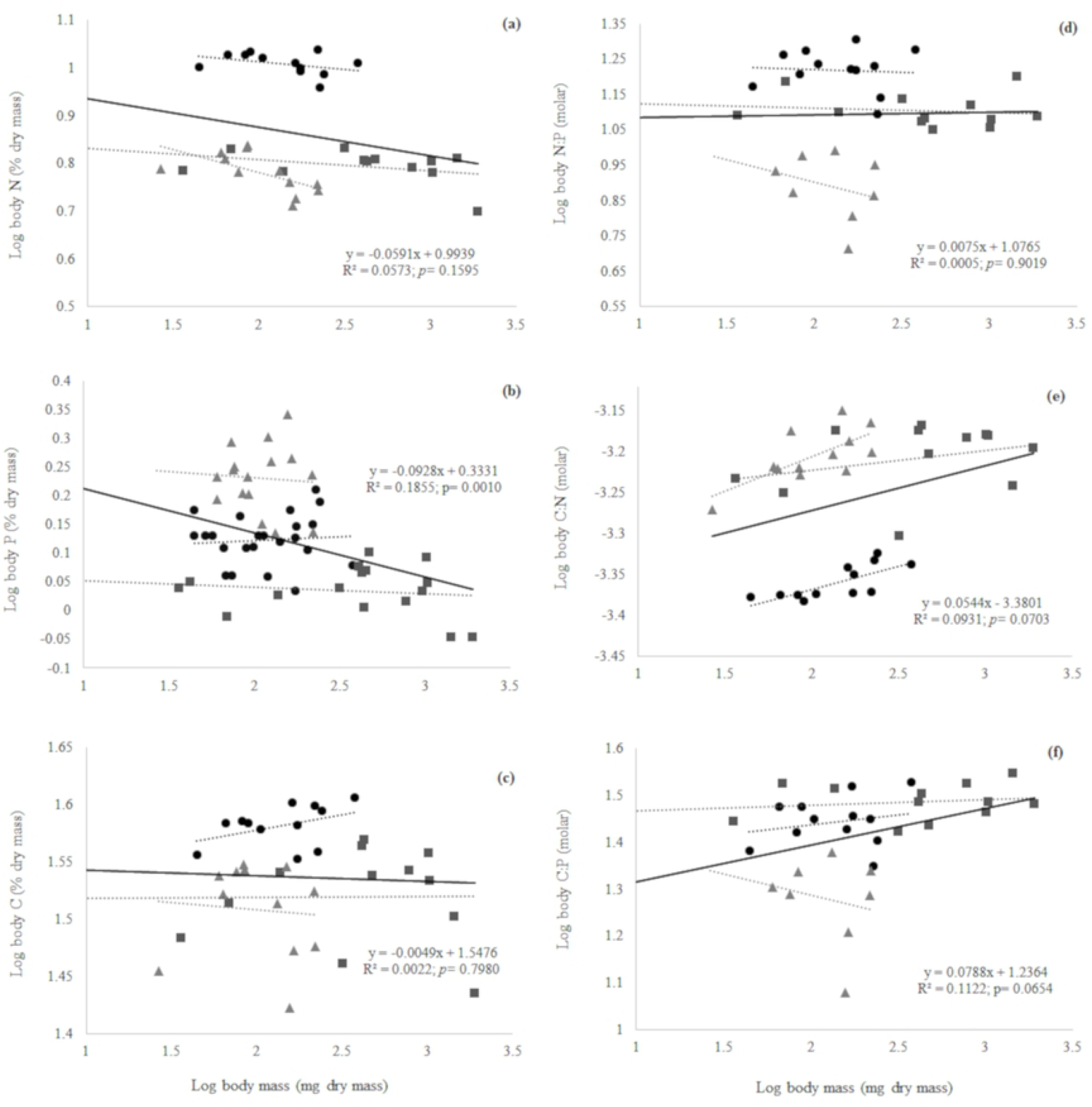
Linear regressions across and within taxa of body content of N, P, C and N: P, C:N, C:P as a function of invertebrate body mass. *Macrobrachium borellii* (black circle), *Aegla uruguayana* (dark gray square), *Trichodactylus borellianus* (light gray triangle).

**Table 5.** Within taxa linear regressions of body elemental content (N, P, C, N:P, C:N, C:P) as a function of decapods’ body mass.

|  | Slope | R<sup>2</sup> | p-value |
| --- | --- | --- | --- |
| Body N |  |  |  |
| *M. borellii* | -0.0311 | 0.1371 | 0.2520 |
| *A. uruguayana* | -0.0232 | 0.1268 | 0.3460 |
| *T. borellianus* | **-0.0957** | **0.3930** | 0.0290 ** |
| Body P |  |  |  |
| *M. borellii* | 0.0133 | 0.0059 | 0.7256 |
| *A. uruguayana* | -0.0115 | 0.0195 | 0.6060 |
| *T. borellianus* | -0.0210 | 0.0041 | 0.8117 |
| Body C |  |  |  |
| *M. borellii* | 0.0265 | 0.1620 | 0.1945 |
| *A. uruguayana* | 0.0009 | 0.0001 | 0.9705 |
| *T. borellianus* | -0.0127 | 0.0067 | 0.7990 |
| Body N:P |  |  |  |
| *M. borellii* | -0.0138 | 0.0038 | 0.8488 |
| *A. uruguayana* | -0.0121 | 0.0180 | 0.6776 |
| *T. borellianus* | -0.1286 | 0.0839 | 0.4865 |
| Body C:N |  |  |  |
| *M. borellii* | **0.0576** | **0.5598** | 0.0051** |
| *A. uruguayana* | 0.0241 | 0.0955 | 0.3284 |
| *T. borellianus* | **0.0845** | **0.4889** | 0.0114 * |

| Body C:P |  |  |  |
| --- | --- | --- | --- |
| *M. borellii* | 0.0437 | 0.0503 | 0.4831 |
| *A. uruguayana* | 0.0119 | 0.0263 | 0.6139 |
| *T. borellianus* | -0.0899 | 0.0413 | 0.6293 |

Statistically significant linear relationships between species body mass and the respective response variable \* $p < 0.05$ , \*\* $p < 0.005$ , \*\*\* $p < 0.001$ .

**b)** Among taxa, it was observed a significant individual effect of the taxonomic identity on body N, P, C, N:P, C:N and C:P contents (ANCOVA, *p*= 2×10^-16^, *p*= 9.56×10^-13^, *p*= 6.84×10^-5^, *p*= 3.95×10^-10^, *p*= 1.05×10^-14^ and *p*= 3.71×10^-7^, respectively) and significant individual effect of body mass on body N (ANCOVA, *p*= 0.0001), body P (ANCOVA, *p*= 3.54×10^-6^) and body C:N contents (ANCOVA, *p*= 0.0052). According to *post hoc* interspecific analysis, body N and C contents were statistically different among *M. borellii* prawn (with highest average values) and the other species (Bonferroni *post hoc*, *p*< 0.001) (Fig 4a, c). This prawn also showed significant differences in body C:N content between the other species (Bonferroni *post hoc*, *p*< 0.001), but with lowest average values (Fig 4e). Percentage of body P and N:P contents were statistically different among all species (Bonferroni *post hoc*, *p*< 0.001) (Fig 4b, d). *T. borellianus* crab showed the highest average value of body P content and *A. uruguayana* aeglid, the lowest one (Fig 4b). Moreover, *M. borellii* prawn showed the highest value of body N:P content and *T. borellianus* crab, the lowest one (Fig 4d). Finally, *T. borellianus* crab showed statistically differences in body C:P content in comparison to the other species (Bonferroni *post hoc*, *p*< 0.001), with the lowest average values (Fig 4f).

**Fig 4.**
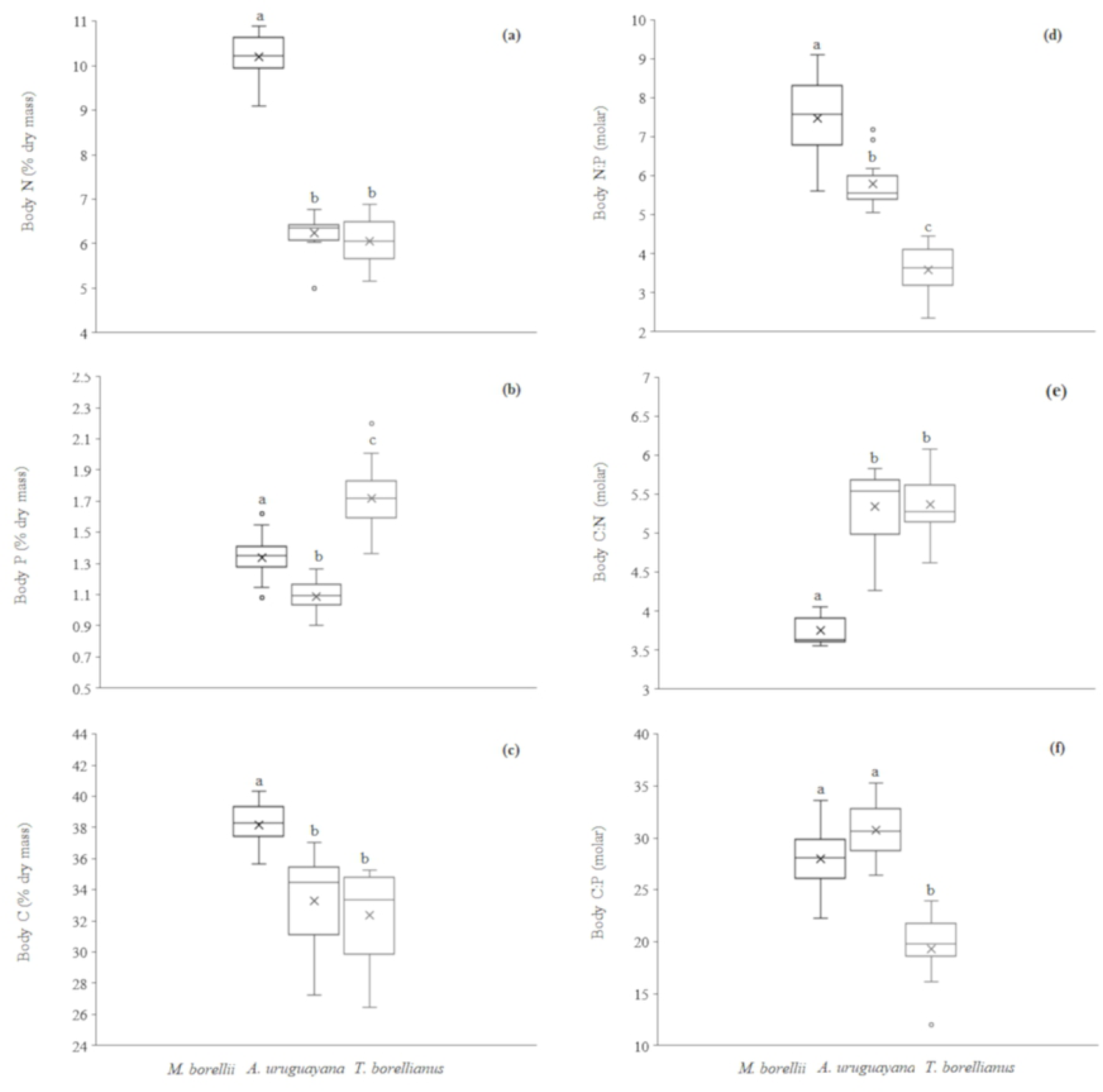
Box plots of body content of N, P, C and N:P, C:N, C:P of each decapod species. The top, bottom, and line through the middle of the boxes correspond to the 75th, 25th, and 50th (median) percentile, respectively. The whiskers extended from the 10th percentile to the 90th percentile. Crosses indicate the median values. Different letters above bars indicate significant differences among taxa (*p* < 0.05).

The first two components of PCA explained the 39.38% (Component 1) and 22.13% (Component 2), showing variations among species (Fig 5). Body N, C, N:P and C:N contents characterised species’ variations with high correlation values in Component 1 (Table 6). Body N, C, and N:P contents were higher in *M. borellii* prawn and lower in the other crustaceans, while body C:N followed the opposite trend. Body P content showed high correlation values in Component 2 (Table 6), being higher in *T. borellianus* and lower in *A. uruguayana* (Fig 5). The body C:P content showed similar values of correlation in both components, and was low in *T. borellianus*. The nutrient excretions and ratio presented low values of correlation in both components. However, it was observed that N excretion was higher in *M. borellii* and lower in *A. uruguayana.* Besides, this variable increased with body N content and decreased with body mass. *T. borellianus* crab exhibited higher P excretion and body P content, which decreased with body mass. The N:P excretion increased in *A. uruguayana* and with body mass. Body mass presented the higher correlation in Component 2, and was higher in *A. uruguayana* (Table 6) (Fig 5).

**Fig 5.**
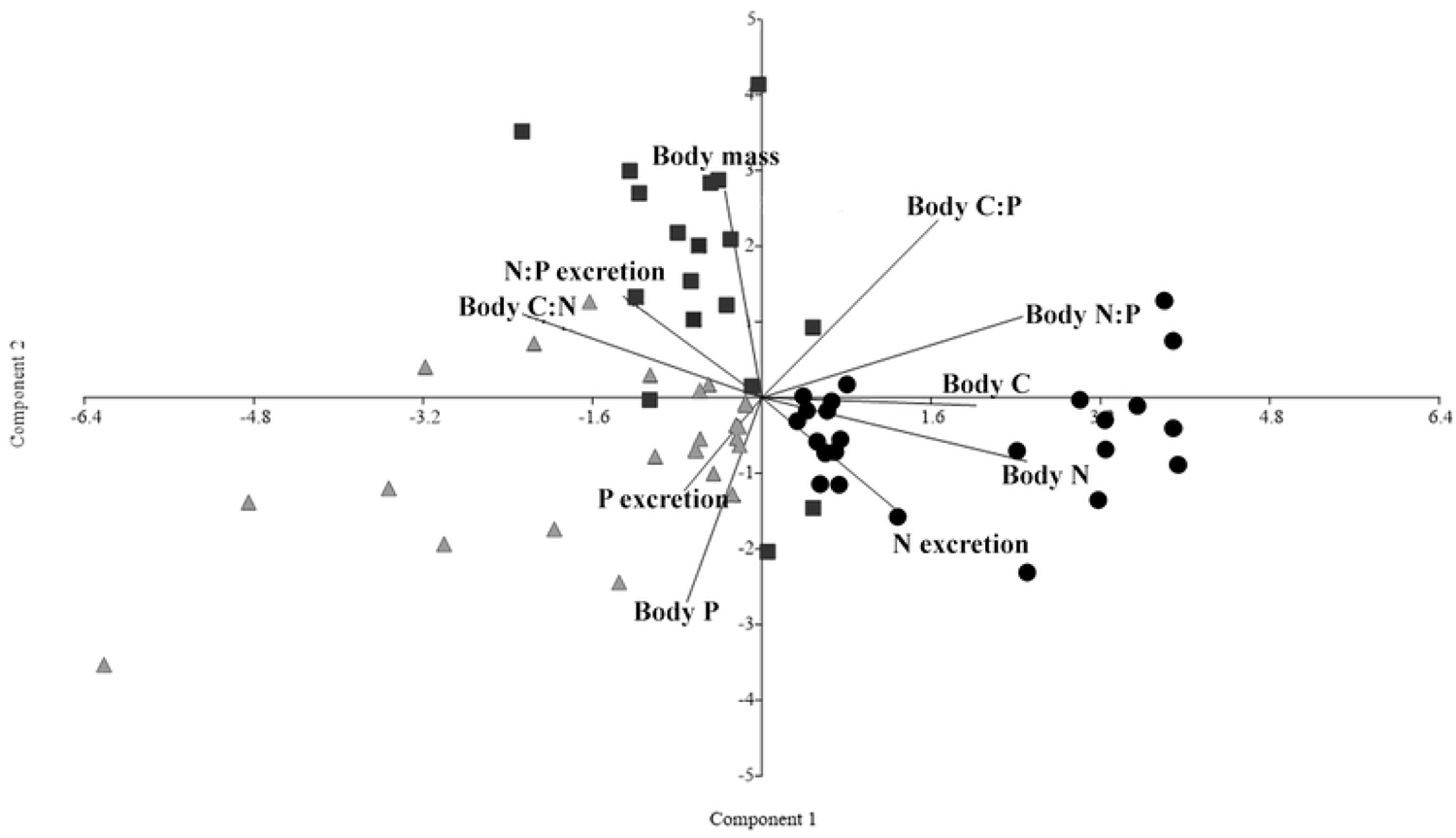
Principal component analysis with log10-transformed of nutrient excretion rates (N and P) and ratio (N:P), body elemental contents (body N, P, C and body N: P, C:N, C:P) and body mass. Arrangement by taxonomy identity of each species. *Macrobrachium borellii* (black circle), *Aegla uruguayana* (dark gray square), *Trichodactylus borellianus* (light gray triangle).

**Table 6.** Correlation values excretion rates (N, P) and ratio (N:P), body nutrient contents (N, P, C) and ratios (N:P, C:N, C:P), and body mass on the first and second principal components of PCA-Principal component analysis.

| Variable | Correlations of PCA |  |
| --- | --- | --- |
|  | Component 1 | Component 2 |
| N excretion | 0.4632 | -0.4048 |
| P excretion | -0.2706 | -0.3407 |
| N:P excretion | -0.4847 | 0.3719 |
| Body N content | 0.9264 | -0.2331 |
| Body P content | -0.262 | -0.749 |
| Body C content | 0.7498 | -0.02835 |
| Body N:P | 0.9116 | 0.2971 |
| Body C:N | -0.8368 | 0.3061 |
| Body C:P | 0.6150 | 0.6495 |
| Body mass | -0.1286 | 0.7587 |

## Discussion

In this study, we assessed which variables best explained nutrient recycling in two scales of analyses (across and within taxa) through empirical data obtained from three freshwater decapod species of different families. Firstly, we analysed the predictive power of physiological (body mass, body elemental content), ecological (diet), taxonomic and experimental (time of incubation) variables on nutrient (N, P, N:P) excretion. Secondly, we focused the analysis on physiological variables to test the hypothesis of ecological theories in our data. We also evaluated and compared the body elemental (C, N, P) content and associated ratios across and within taxa, and used physiological variables as covariates among taxa.

**Q1-** Our results showed that the predictive power of variables analysed together changed with the scale of analyses, but the importance of body mass and taxonomic identity on nutrient excretion rates were observed across and within taxa **(H1)**. The time of incubation was also a factor that should be considered in experimental design and, in our case, 30 minutes was the most recommended one.

**Q2-** When physiological variables (body mass, body elemental content) were analysed independently, body size was a better predictor of nutrient mineralization than body elemental content in both scales of analyses. N:P excretion scaled negatively with body N:P content only across taxa **(H2)**. Allometry was observed across and within taxa, mainly for N excretion; that is, juveniles mineralized more nutrients than adults **(H3)**. Among taxa, the taxonomic identity and body mass had a strong influence in the nutrient N and P excretion while taxonomic identity and body N content had a significant effect on N excretion **(H4)**.

**Q3-** Regarding the body elemental content of crustaceans, only body P content of decapods scaled negatively with body mass across taxa **(H5)**, while within taxa, the body N of crabs (negative slope) and body C:N of prawns and crabs (positive slope) exhibited allometry. Among taxa, all elements and ratios were different and a significant effect of body mass was observed on body N, P and C:N. Body C:N was higher in carcinized decapods in comparison with prawns, while body N:P was different in all species and higher in prawns, species with more carnivorous trophic habit **(H6)**. We discussed the implications of the main results, their relations with ecological theories and the usefulness in productive aquatic systems such as IMTA.

**Q1** *-* Several studies have shown that body size (by extension, body mass) is the best predictor of N and P excretion rates in fish, decapod crustaceans and other invertebrates [18,20,22,38,100,101]. This relationship is even more marked in the excretion of N than in P [6], such as observed in the present study. The results obtained by [20] and [6] highlighted the primary importance of MTE in predicting nutrient flux through organisms over other predictors, such as the body elemental composition and diet. Physiological variables like body mass are unfailingly related to taxonomic identity, as corroborated in the present study. For example, the rate at which nutrients are assigned to processes such as reproduction, growth and feeding events are conserved and characteristic at a certain taxonomic level [3, 20]. In this way, it is expected that differences observed within species must be due to specific features of life cycle and trophic habits [47, 54] of prawn, anomuran and crab species studied here.

We also noticed changes in excretion rates over time in both scales of analyses. The nutrient excretion rate was higher during the first 30 minutes of incubation and decreased at 60 minutes. This difference was significant for N excretion across taxa and in *A. uruguayana* within taxa. Previous studies also reported reduction of nutrient excretion rates as regards time and, in general, authors attributed the reduction in mineralization rates with time due to fasting and handling stress [15,21,22,92]. In our case, crustaceans were acclimated to experimental conditions during 2 weeks, and fed the same diet during this period and just before the excretion trial. In this way, we could infer that differences found in N excretion time had low bias due to fasting and handling stress, and that 30 minutes was an adequate incubation period to make comparative analyses with the studied crustaceans. [21] and [15] also recommended this time lapse as the most appropriate minimum period for the excretion rates to stabilize. The significantly predictive power of time on N excretion rate, which was not observed in P excretion, could be due to the animals’ necessity to rapidly remove toxic ammonia from protein metabolism, which negatively influences the organisms’ homeostasis [102].

The other evaluated variables –diet and body composition– had no effect on the variations in nutrient excretion rates across taxa. Within taxa, only body N content of *M. borelli* positively affected N excretion. Regarding diets used in excretion trials, they had formulations for omnivorous and detritivorous fish that did not differ in N and P content, allowing the two groups of data to be analysed together. Additionally, crustaceans fed the same diet for 2 weeks, so it was expected that excretion rates were not influenced by the variable trophic resource, as it commonly occurs in field experiments [103]. Therefore, it was expected that changes in nutrient excretion were related to body elemental composition, which was not true either.

The importance of diet and body elemental composition in predicting excretion rates are key concepts in EST, but our results showed that these variables have little or no predictive power to our data. [15] and [20] also found that diet and body elemental content were not good predictors of excretion rates. [17] concluded that diet was not a good predictor, and both N and P excretion rates did not reflect the trophic position (determined by δ^15^N signatures) of shrimp and fish species. In the case of *M. borellii*, within taxa analyses showed that the body N content of this prawn positively affected N excretion rate, contrary to the EST predictions. However, this regression was not linear and results should be corroborated with excretion trials that evaluate nutrient excretion rates against the consumption of diets with variable N and P composition.

**Q2** - The independent analyses of physiological variables supported body mass as a better predictor of nutrient mineralization than body elemental content, both across and within taxa, emphasizing the results found for question 1 (Q1). As previously predicted by MTE, the relationship between these variables is negative [3,7,10] and exhibited variable slopes and explanation percentage at the temperature (24 ± 1 °C) used in the experiments. While body mass explained 45% and 13% of variation in the excretion rates of N and P across taxa, respectively, body N content only explained 36% of N excretion and exhibited a positive slope, which contradicts the EST predictions. On the contrary, body N:P content explained 18% of N:P excretion with a negative slope, fulfilling what was predicted by the EST, despite the low explanation. When we explored results within taxa, all species presented a negative and significant relation between N excretion and body mass, and the regression for *A. uruguayana* presented higher explanation (85%) than the other crustaceans. In this species, body mass also explained 65% of the P excretion rates.

The greater predictive power of the body mass in comparison to body elemental content is in concordance with previous studies [6, 20] that found little support for the predictions of EST in aquatic organisms. This is reasonable since the excretion of waste nutrients is related to the acquisition and assimilation of food, both processes driven by metabolic demands that change during the life cycle of the organisms [6, 7]. Although several studies highlighted the effect of body elemental content on excretion rates in various groups of animals [13, 38], for the analysed crustacean species, this trait has a weak predictive value [6,18,20].

Although MTE predictors (body mass) explained more variance than EST predictors (body elemental content), the scaling coefficients (regression slopes) revealed variable values in both scales of analysis. Across taxa, slopes were −0.68 for N and −0.49 for P, showing a steeper decrease of N excretion rates than P as crustaceans grew. Within taxa, significant values ranged from −0.56 to −0.75 for N, and −0.67 for P in *A. uruguayana*. In this case, only anomurans exhibited significant relation between both N and P excretion and body mass with higher slope values than the other species. These results indicated that juveniles of the three crustaceans species mineralized more N than adults, and that only juveniles of *A. uruguayana* mineralized more P than adults. The excretion rate per unit mass of juveniles, which is higher than adults (negative slope), was also observed in other crustaceans, mainly for N [18,22,38]. The steeper or shallower scaling of excretion rates observed in the three crustacean species, and under the consumption of the same diet, is complex to explain only from an MTE perspective [6]. Body mass and taxonomic identity are not independent, and determining the relative contribution of each one in the variation of excretion rates is a challenge [10].

The analyses performed among taxa emphasized that the taxonomic identity had a strong influence in the nutrient excretion rates using both body mass and body elemental content as covariates. These last variables influenced N excretion, while in body elemental content, they only influenced P excretion. In prawns, body N content and body mass influenced the N excretion, differing from crabs and anomurans. Contrary to what would be expected from the EST predictions, prawns mineralized higher N than the other species, even with higher body N content and after the consumption of diets that did not fulfill the protein requirement of this species to grow (∼35%) [60]. *A. uruguayana* anomuran and *T. borellianus* crab released the intermediate and the lowest rates of N, respectively, showing the same trend in their respective amounts of body N; results that do not either align with EST predictions [8]. These results are surprising and, as suggested by [104] in armour fish taxa, could be related to the variation in the assimilation efficiency, and even in variations in the phenotype expression given the high individual variability observed mainly in the N excretion of prawns.

Future studies should include egestion rates of these organisms and feed digestibility to help understand differences in assimilation efficiency in each taxon [105]. These feeding trials would also be helpful to quantify elemental imbalances between body and diet, and to verify the homeostasis regulation of these species [8]. Such trials should consider the residence time and turnover of elements in crustaceans’ tissues [106] in the experimental design. Regarding P excretion rates, *T. borellianus* excreted higher amounts than *M. borellii* (lower values) and *A. uruguayana.* In heterotrophic organisms, the assimilated P is stored as polyphosphate rich in energy [107, 108]. Therefore, prawns and anomurans would assimilate a greater amount of this nutrient because they are more active than crabs and might have higher energy expenditure in mobility.

Despite the complexity to understand the differences observed in excretion rates among species and the need to complement the results with additional studies, our findings emphasize the importance of the taxonomic identity and the resolution to analyse animal-mediated nutrient cycling [10]. The digestive morphology and physiology are useful tools that play an important role in the excretion process and that could aid understanding the variations in the studied species and during ontogeny. Juveniles of *T. borellianus* and *A. uruguayana* ingest more animal items than adults [55,58,109], while *M. borellii* seem to maintain a preference to animal components during their entire life cycle [54]. Morphological traits of chelae and mandibles of *A. uruguayana* are different between juveniles and adults [109] and, in *T. borellianus,* feeding apparatus are adapted to cope with dietary changes [63, 110]. As regards digestive physiology, juveniles of *A. uruguayana* exhibited higher amylase activity than adults [68] and the metabolic profile varied with the season and ontogeny [67]. In *M. borellii*, the proteinase activity is smaller in juveniles than in adults [68].

The mentioned background information, together with the results of the present study, provides a deeper insight into feeding strategies, nutritional requirements and resource utilization. This allows a thorough comprehension of the role of the studied crustaceans in the cycling of nutrients in natural aquatic systems, in order to estimate the bioremediation efficiency of decapods in IMTA systems, in later studies.

**Q3-** Across taxa analysis showed that crustaceans exhibited a negative relationship between the body P content and body mass with low explanation (18%). The fact that, at order scale, crustaceans seem to follow the GRH hypothesis –which proposes that small-bodied organisms tend to have faster growth rates and higher body P allocation to protein synthesis [8, 11] than larger organisms–, the effect was not observed within taxa. This might have occurred because, when all species are analysed together, mass range is wider and sufficient to observe a difference in body P allocation between organisms of lower and higher mass; that is, low body mass species have higher P content and vice versa. This explanation also makes sense if we consider that the slope of regressions within taxa is close to zero; that is, if there is no difference in the body P content between juveniles and adults within taxa, the relationship between body mass and body P content across taxa is an effect of the scale of analyses and must be carefully interpreted. Body mass also poorly explains animal body elemental content in other studies [23, 25]. The relationship among these variables is not simple and results found in literature are quite variable in across or within taxa analysis, and even more if data is controlled for phylogenetic effects [23,24,26,111].

In our study, within taxa analysis revealed that body N content scaled with body mass in *T. borellianus*. Adult crabs had less N than juveniles and this finding could be related to the relative importance of animal items in the natural diet of this species. Larger individuals consume more vegetal remains than smaller ones, which consume more insect larvae and oligochaetes [56, 90]. This feeding habit is also in congruence with the positive allometry of body C:N observed in this crab, since adults eat more C-rich items. We also observed a higher body C:N in adults of *M. borellii*. However, prawns seem to maintain a preference for animal components during the entire life cycle, contradicting this allometric tendency of body C:N content. If there is no evidence that diet could drive variation in body stoichiometry during ontogeny, this relationship could be related to differences in the proportion of chitin between juveniles and adults. Since the chitin of exoskeleton has high C:nutrient [8,27,112–114], adults of both crabs and prawns should have higher amounts of C than N due to more robust carapace. It is necessary to complement these results with studies that explore the influence of diet on body stoichiometry through ontogeny and the deviation from strict homeostasis to get a better understanding of nutrient allocation strategies. We also recommend a complementary study with a broad range of body mass of each species (mainly for *T. borellianus*) to confirm the allometric trends of body elemental content found in our study.

Regarding among taxa analysis, crustacean species exhibited different body stoichiometry of C, N, P and associated ratios. Prawns had higher body N and C contents, and also lower C:N than anomurans and crabs. These findings could be related to the natural diet and the carnization of crustaceans, since *M. borellii* have a more carnivorous trophic habit [54, 57]. This species requires high protein content to achieve better growth parameters [60] and it is an uncarcinized decapod. These traits could contribute to a lower body C:N content compared to the other species, due to a higher ingestion of N-rich food items (higher trophic status) and to a less chitinous carapace. Moreover, digestive enzyme studies revealed that *M. borellii* had higher proteinase activity while juveniles of *A. uruguayana* had higher amylase activity in hepatopancreas [68], indicating that metabolism of nitrogen and carbohydrates, respectively, predominates in these organisms. The not intuitive high body C content of *M. borellii* could be due to the fact that crustaceans store more lipids in muscle tissue than anomurans [68] and that they also have comparatively more muscle tissue (therefore more body N), which might explain lower values of body C:N. On the other hand, crabs-like body forms species (robust carapace with lower N-rich bodies), such as *T. borellianus* and *A. uruguayana*, also consume more algae and vegetal remains than prawns [54,58,106]. The lower trophic status of crabs and anomurans, therefore, exhibited more C:N as expected.

Further, body P and N:P content were different among the three species, with *T. borellianus* presenting the higher and lower values, respectively. This crab seems to have higher specific growth rates than prawns and anomurans when fed the same omnivorous feed used in this study (Calvo et al., unpublished data). Therefore, higher body P content should be related with a greater allocation of this element to the production of P-rich rRNA to hold faster growth [4] and the lowest N:P ratio is explained by the high P content with respect to N in this species’ body. However, this hypothesis should be corroborated by carrying out growth experiments that ensure there is no deficiency in any element regarding the feed formulation, and that it matches the nutritional requirements of tested species.

### Comments about application in IMTA

Aquaculture represents almost 50% of the fishery products produced globally and continues to grow faster than other food production sectors, mainly in freshwater systems [115]. The nutrient discharge of this food production system has generated many environmental impacts on ecosystems [116] that could be mitigated and recycled through the diversification and incorporation of complementary species, such as proposed by IMTA concepts [117, 118]. In low taxa diversity aquatic systems, like an IMTA, the nutrient turnover is strongly influenced by the few taxa that compose it [81, 14]. Therefore, understanding the physiological processes of organisms at different levels of organizations is essential to explore and predict nutrient recycling behavior in different scenarios (unique species and groups of species) and to design integrated productive systems that efficiently use the nutrient inputs through an adjusted mass balance.

Decapods could display different roles as organic nutrient recyclers in integration with detritivorous and omnivorous fish. The differences found in N and P excretion rates across and within taxa highlighted that if they were integrated as a unique species or as a group of species, the effect on nutrient recycling could be different. In both cases, we expected that juveniles recycle more nutrients per mass unity than adults, mainly for N. Therefore, early stage decapods would contribute more to transforming N from proteins to ammonium (i.e. reducing organic burden) and enhance the availability of this compound to nitrifying bacteria to finally grow vegetables or algae. Inorganic N could be even more available with the integration of prawns only. On the other hand, prawns incorporate more P and, therefore, could limit this nutrient to primary producers while representing a P-rich by-product for many purposes. Contrary, crabs mineralize less N and more P exerting the opposite effect. These results showed that the role of crustaceans as animal-mediate nutrient recyclers depends mainly on body mass and taxonomic identity and should be considered to select complementary species that efficiently use feed resources.

Body elemental content is another variable that could be a useful baseline to estimate the nutritional requirements of animals [119, 120]. Our study revealed that this variable differed with body mass (ontogeny) and that it is related to the natural trophic status of organisms. Results showed higher N-requirements and higher trophic status (lower C:N) of prawns than anomurans and crabs. Moreover, prawns and crabs change the trophic status during ontogeny, requiring lower N in comparison with C at later stages. Prawns need more protein in the feed and might be successfully integrated with fish of higher N-requirements, while crabs and anomurans might exhibit good performance with fish that are fed lower N-rich diets, such as herbivorous and omnivorous fish.

Our study was an effort to contribute to the ecological background of MTE and EST through empirical data obtained from freshwater decapod species with aquacultural potential use, and to provide useful information about nutrient mineralization and nutritional requirements based on body elemental content of these crustaceans. These results, added to other empirical data on egestion, digestibility, retention in biomass (growth and reproduction), and food intake, offer a framework to leave some open questions for further studies and provides information to infer the amount of crustaceans needed to biomitigate the feed remains of detritivorous and omnivorous fish culture, through an improved mass balance. Ecological theories and experimental aquaculture research can be good allies when it is necessary to “imitate” nature and achieve more efficient production processes with less impact on the ecosystem.

## Acknowledgments

We thank C. De Bonis for his field assistance and M. C. Mora for her lab assistance.

## References

1. Vanni MJ, Boros G, McIntyre PB. When are fish sources vs. sink of nutrients in lake ecosystems? Ecology. 2013; 94(10): 2195–206.

2. Atkinson CL, Capps KA, Rugenski AT, Vanni MJ. Consumer-driven nutrient dynamics in freshwater ecosystems : from individuals to ecosystems. Biol Rev. 2017; 92(4): 2003–23.

3. Vanni MJ. Nutrient cycling by animals in freshwater ecosystems. Annu Rev Ecol Syst. 2002; 33: 341–70.

4. Elser JJ, Acharya K, Kyle M, Cotner J, Makino W, Markow T, Watts T, Hobbie S, Fagan W, Schade J, Hood J, Sterner RW. Growth rate – stoichiometry couplings in diverse biota. Ecol Lett. 2003; 936–43.,

5. Karasov WH, Martinez del Río C. Physiological Ecology: How animals process energy, nutrients, and toxins. Princeton, 2007.

6. Vanni MJ, Mcintyre PB. Predicting nutrient excretion of aquatic animals with metabolic ecology and ecological stoichiometry : a global synthesis Author (s): Michael J. Vanni and Peter B. McIntyre Published by : Wiley on behalf of the Ecological Society of America Stable URL. 2016; 97(12): 3460–3471.

7. Brown JH, Gillooly JF, Allen AP, Savage VM, West GB. Toward a metabolic theory of ecology. Ecology. 2004; 85(7): 1771–1789.

8. Sterner RW, Elser JJ. Ecological Stoichiometry. The biology of elements from molecules to the biosphere. Princeton University Press.; 2002.

9. Woodward G, Ebenman B, Emmerson M, Montoya JM, Olesen JM, Valido A, et al. Body size in ecological networks. 2005; 20(7): 402–409.

10. Hall RO, Koch BJ, Marshall MC, Taylor BW, Tronstad LM. How body size mediates the role of animals in nutrient cycling. In: Hildrew AG, Raffaelli DG, Edmonds-Brown R, editors. Body Size: The Structure and Function of Aquatic Ecosystems. Cambridge University Press, Cambridge; 2007.

11. Elser JJ, Sterner RW, Gorokhova E, Fagan WF, Markov TA, Cotner JB, et al. Biological stoichiometry from genes to ecosystems. Ecol Lett. 2000; 3: 540–50.

12. Sterner RW. The ratio of nitrogen to phosphorus resupplied by herbivores: zooplankton and the algal competitive arena. Am Nat. 1990; 136(2): 209–29.

13. Vanni MJ, Flecker S, Hood JM. Stoichiometry of nutrient recycling by vertebrates in a tropical stream : linking species identity and ecosystem processes. Ecol Lett. 2002; 5(2): 285–93.

14. Torres LE, Vanni MJ. Stoichiometry of nutrient excretion by fish: interspecific variation in a hypereutrophic lake. Oikos. 2007; 116: 259–270.

15. Mcmanamay RA, Webster JR, Valett HM, Dolloff CA, Mcmanamay RA, Webster JR, et al. Does diet influence consumer nutrient cycling? Macroinvertebrate and fish excretion in streams. J North Am Benthol Soc. 2011; 30(1): 84–102.

16. Dalton CM, El-sabaawi RW, Honeyfield DC, Auer SK, Reznick N, Flecker AS. The influence of dietary and whole-body nutrient content on the excretion of a vertebrate consumer. PLoS ONE. 2017; 12(11): e0187931.

17. Zandonà E, Oliveira-cunha P, Tromboni F, Neres-lima V, Moraes M, Moulton TP. Do body elemental content and diet predict excretion rates of fish and shrimp? Fundam Appl Limnol. 2021; 3: 271–83.

18. Benstead JP, Cross WF, March JG, Mc Dowell WH, Ramírez A, Covich AP. Biotic and abiotic controls on the ecosystem significance of consumer excretion in two contrasting tropical streams. Freshw Biol. 2010; 55(10): 2047–61.

19. Moody EK, Corman JR, Elser JJ, Sabo JL. Diet composition affects the rate and N : P ratio of fish excretion. Freshw Biol. 2015; 60(3): 456–65.

20. Allgeier JE, Wenger SJ, Rosemond AD, Schindler DE, Layman C. Metabolic theory and taxonomic identity predict nutrient recycling in a diverse food web. Proc Natl Acad Sci U S A. 2015; 112: e2640–e2647.

21. Whiles MR, Huryn AD, Taylor BW, Reeve JD. Influence of handling stress and fasting on estimates of ammonium excretion by tadpoles and fish : recommendations for designing excretion experiments. Limnol Oceanogr Methods. 2009; 7(1): 1–7.

22. Oliveira-cunha P, Capps KA, Lourenc C, Tromboni F, Moulton TP. Effects of incubation conditions on nutrient mineralisation rates in fish and shrimp. Freshw Biol. 2018; 63: 1107–1117.

23. González AL, Fariña JM, Kay AD, Pinto R, Marquet PA. Exploring patterns and mechanisms of interspecifi c and intraspecifi c variation in body elemental composition of desert consumers. Oikos. 2011; 120: 1247–55.

24. González AL, Farjalla VF, Dézerald O, Leroy C, Richardson BA, Romero GQ, et al. Ecological mechanisms and phylogeny shape invertebrate stoichiometry : A test using detritus - based communities across Central and South America. Funct Ecol. 2018; 32: 2448–63.

25. Allgeier J, Wenger S, Layman A. Taxonomic identity best explains variation in body nutrient stoichiometry in a diverse marine animal community. Sci Rep. 2020; 10: 13718.

26. Andrieux B, Signor J, Guillou V, Danger M. Body stoichiometry of heterotrophs : Assessing drivers of interspecific variations in elemental composition. Glob Ecol Biogeogr. 2021; 30(4): 883–95.

27. Elser JJ, Dobberfuhl DR, Mackay NA, Schampel JH. Organism size, life history, and N:P stoichiometry. Bioscience. 1996; 46(9): 674–84.

28. Faerøvig PJ, Hessen DO. Allocation strategies in crustacean stoichiometry: the potential role of phosphorus in the limitation of reproduction. Freshw Biol. 2003; 48: 1782–92.

29. Zhang P, van den Berg RF, van Leeuwen CHA, Blonk BA, Bakker ES. Aquatic omnivores shift their trophic position towards increased plant consumption as plant stoichiometry becomes more similar to their body stoichiometry. PLoS One. 2018; 13: 1–13.

30. Benstead JP, Hood JM, Whelan NV, Kendrick MR, Nelson D, Hanninen AF, et al. Coupling of dietary phosphorus and growth across diverse fish taxa: a meta-analysis of experimental aquaculture studies. Ecology. 2014; 95(10): 2768–77.

31. Frost PC, Cross WF, Benstead JP. Ecological stoichiometry in freshwater benthic ecosystems : an introduction. Freshw Biol. 2005; 50(11): 1781–5.

32. Pilati A, Vanni MJ. Ontogeny, diet shifts, and nutrient stoichiometry in fish. Oikos. 2007; 116: 1663–74.

33. McIntyre PB, Flecker AS, Vanni MJ, Hood JM, Taylor BW, Thomas SA. Fish distributions and nutrient cycling in streams: can fish create biogeochemical hotspots? Ecology. 2008; 89(8): 2335–46.

34. Christian AD, Crump BG, Berg DJ. Nutrient release and ecological stoichiometry of freshwater mussels (Mollusca: Unionidae) in 2 small, regionally distinct streams. J North Am Benthol Soc. 2008; 27(2): 440–50.

35. Atkinson CL, Opsahl SP, Covich AP, Golladay SW, Conner LM. Stable isotopic signatures, tissue stoichiometry, and nutrient cycling (C and N) of native and invasive freshwater bivalves. J North Am Benthol Soc. 2010; 29(2): 496–505.

36. Evans-White MA, Lamberti GA. Grazer species effects on epilithon nutrient composition. Freshw Biol. 2005; 50: 1853–63.

37. Cross WF, Covich AP, Crowl TA, Benstead JP, Ramírez A. Secondary production, longevity and resource consumption rates of freshwater shrimps in two tropical streams with contrasting geomorphology and food web structure. Freshw Biol. 2008; 53: 2504–19.

38. Alves JM, Caliman A, Guariento RD, Figueiredo-Barros MP, Carneiro LS, Farjalla VF, et al. Stoichiometry of benthic invertebrate nutrient recycling : Interspecific variation and the role of body mass interspecific variation and the role of body mass. Aquat Ecol. 2010; 44(2): 421–30.

39. Tsoi WY, Hadwen WL, Fellows CS. Spatial and temporal variation in the ecological stoichiometry of aquatic organisms in an urban catchment. J North Am Benthol Soc. 2011; 30: 533–45.

40. Snyder MN, Small GE, Pringle CM. Diet-switching by omnivorous freshwater shrimp diminishes differences in nutrient recycling rates and body stoichiometry across a food quality gradient. Freshw Biol. 2015; 60(3): 526–36.

41. Pringle CM, Hamazaki T. The role of omnivory in a neotropical stream: separating diurnal and nocturnal effects. Ecology. 1998; 79: 269–80.

42. Crowl TA, McDowell WH, Covich AP, Johnson S. Freshwater shrimp effects on detrital processing and nutrients in a tropical headwater stream. Ecology. 2001; 82: 775–783.

43. Dobson M, Magana A, Mathooko JM, Ndegwa FK. Detritivores in Kenya Highland streams: more evidence for the paucity of shredders in the tropics? Freshw Biol. 2002; 47: 909–19.

44. Cogo GB, Santos S. The role of aeglids in shredding organic matter in neotropical streams. J Crustac Biol. 2013; 33: 519–26.

45. Melo GAS. Manual de identificação dos crustáceos decápodos de água doce do Brasil. São Paulo: Editora Loyola; 2002.

46. Collins PA, Giri F, Williner V. Population dynamics of Trichodactylus borellianus (Crustacea Decapoda Brachyura) and interactions with the aquatic vegetation of the Paraná River (South America, Argentina). Ann Limnol J Limnol. 2006; 42: 19–25.

47. Collins PA, Williner V, Giri F. Littoral communities. Macrocrustaceans. In: Iriondo MH, Paggi JC, Parma MJ, editors. The Middle Paraná River: Limnology of a Subtropical Wetland. Springer-Verlag; 2007.

48. Williner V, Giri F, Collins PA. Ontogeny of the mandible of Aegla uruguayana: A geometric morphometric approach. Paleontol I Evol Mem Espec. 2009; 3: 117–8.

49. Bond-Buckup G, Jara CG, Pérez-Losada M, Buckup L, Crandall KA. Global diversity of crabs (Aeglidae: Anomura: Decapoda) in freshwater. Hydrobiologia. 2008; 595: 267–273.

50. Crandall KC, Buhay JE. Global diversity of crayfish (Astacidae, Cambaridae, and Parasticidae-Decapoda) in freshwater. Hydrobiologia. 2008; 595: 295–301.

51. De Grave S, Cai Y, Anker A. Global diversity of shrimps (Crustacea: Decapoda: Caridea) in freshwater. In: Balian EV, Lévêque C, Segers H, Martens K, editors. Freshwater Animal Diversity Assessment. Springer; 2008.

52. Yeo DCJ, Peter KL, Cumberlidge N, Magalhães C, Daniels SR, Campos MR. Global diversity of crabs (Crustacea: Decapoda: Brachyura) in freshwater. Hydrobiologia. 2008; 595: 275–286.

53. Scholtz G, Berlin H, Biologie I, Zoologie V. Evolution of crabs – history and deconstruction of a prime example of convergence. Contrib to Zool. 2014;83(2):87– 105.

54. Collins PA, Williner V, Giri F. Trophic relationships in Crustacea Decapoda of a river with floodplain. In: Elewa AMT, editor. Predation in Organisms: A Distinct phenomenon. Springer, Verlag; 2007.

55. Williner V. Foregut ossicles morphology and feeding of the freshwater anomuran crab Aegla uruguayana (Decapoda, Aeglidae). Acta Zool. 2010; 91(4): 408–415.

56. Carvalho DA, Williner V, Giri F, Vaccari C, Collins P. Quantitative food webs and invertebrate assemblages of a large River : a spatiotemporal approach in floodplain shallow lakes. Mar Freshw Res. 2016; 68: 293–307.

57. Collins PA, Paggi JC. Feeding ecology of Macrobrachium borellii (Nobili) (Decapoda: Palaemonidae) in the flood valley of the River Paraná, Argentina. Hydrobiologia. 1997; 362: 21–30.

58. Williner V, Collins PA. Feeding ecology of the freshwater crab Trichodactylus borellianus (Decapoda: Trichodactylidae) in the floodplain of the Paraná River, southern South America. Lat Am J Aquat Res. 2013; 41: 781–92.

59. Williner V, Carvalho D, Collins PA. Feeding spectra and activity of the freshwater crab Trichodactylus kensleyi (Decapoda: Brachyura: Trichodactylidae) at La Plata basin. Zool Stud. 2014; 53(1): 71.

60. Carvalho DA, Reyes PS, Williner V, Mora MC, De Bonis CJ, Collins PA. Growth, survival, body composition and amino acid profile of *Macrobrachium borellii* against the limitation of feeds with different C:N ratios with comments about application in integrated multi-trophic aquaculture. Aquac Res. 2020; 51(10): 3947–3858.

61. Musin GE, Carvalho DA., Viozzi MF, Mora MC, Collins PA, Williner V. Protein and cellulose level in diet: effects on enzymatic activity, metabolite and amino acid profiles in freshwater anomurans *Aegla uruguayana* (Decapoda: Anomura). Aquac Res. 2020; 51(3): 1232–1243.

62. Collins PA. Mecanismos de coexistencia en poblaciones de palaemónidos dulciacuícolas. PhD Thesis, Universidad Nacional de La Plata, Argentina. 2000

63. Carvalho DA, Viozzi MF, Collins PA, Williner V. Functional morphology of comminuting feeding structures of *Trichodactylus borellianus* (Brachyura, Decapoda, Trichodactylidae), an omnivorous freshwater crab. Arthropod Struct Dev. 2017; 46:472–482.

64. Carvalho DA, Collins PA. Predation ability of freshwater crabs: age and prey-specific differences in Trichodactylus borellianus (Brachyura: Trichodactylidae). J Freshw Ecol. 2013; 4: 573–584.

65. Carvalho DA, Collins PA, De Bonis C. Ontogenetic predation capacity of *Macrobrachium borellii* (Caridea: Palaemonidae) on prey from littoral-benthic communitie. Nauplius. 2011, 19, 71–77.

66. Gutierrez MF, Rojas Molinas F, Carvalho DA. Behavioral responses of freshwater zooplankton vary according to the different alarm signals of their invertebrate predators. Mar Freshwa Behav Physiol. 2012; 45(5): 317–331.

67. Musin GE, Rossi A, Diawol V, Collins PA, Williner V. Dynamic metabolic pattern of Aegla uruguayana (Schmitt, 1942) (Decapoda: Anomura: Aeglidae): responses to seasonality and ontogeny in a temperate freshwater environment. J Crust Biol. 2017; 37(4): 436-444.

68. Musin G, Rossi A, Diawol VP, Collins PA, Williner V. Development of enzymes during ontogeny of two freshwater Decapoda: *Aegla uruguayana* (Aeglidae) and *Macrobrachium borellii* (Palaemonidae). Aquac Res. 2018; 49: 3889–3897.

69. Diawol V, Torres MV, Collins PA. Field evaluation of oxygen consumption by two freshwater decapod morphotypes (Trichodactylidae and Aeglidae); the effect of different times of the day, body weight and sex Mar Freshwa Behav Physiol. 2016; 49(4): 251–263.

70. Ross LG, Martinez-Palacios CA, Morales EJ. Developing native fish species for aquaculture: The interacting demands of biodiversity, sustainable aquaculture and livelihoods. Aquac Res. 2008; 39(7): 675–683.

71. Somoza GM, Ross LG. Introduction to the special issue on “Development of native species for aquaculture in Latin America II”. Aquac Res. 2011; 42(6): 737–897.

72. García-Guerrero M, Santos Romero R, Vega-Villasante F, Cortes-Jacinto E. Conservation and aquaculture of native freshwater prawns: the case of the cauque river prawn Macrobrachium americanum (bate, 1868). Lat Am J Aquat Res. 2015; 43(5): 819-827.

73. David FS, Proença DC, Valenti WC. Nitrogen budget in integrated aquaculture systems with Nile tilapia and Amazon River prawn. Aquac Int. 2017; 25: 1733–1746. https://doi.org/10.1007/s10499-017-0145-y

74. David FS, Proença DC, Valenti WC. Phosphorus budget in integrated multi-trophic aquaculture systems with Nile tilapia, *Oreochromis niloticus*, and Amazon River prawn, Macrobrachium amazonicum. JWAS. 2017; 48: 402–414.

75. Dantas DP, Flickinger DL, Costa GA, Batlouni SR, Moraes-Valenti P, Valenti WC. Technical feasibility of integrating Amazon river prawn culture during the first phase of tambaqui growout in stagnant ponds, using nutrient-rich water. Aquac. 2020; 516: 734611

76. Flickinger DL, Costa GA, Dantas DP, Moraes-Valenti P, Valenti WC. The budget of nitrogen in the grow-out of the Amazon river prawn (*Macrobrachium amazonicum* Heller) and tambaqui (*Colossoma macropomum* Cuvier) farmed in monoculture and in integrated multi-trophic aquaculture systems. Aquac Res. 2019; 50: 3444–3461.

77. Flickinger DL, Dantas DP, Proença DC, Valenti WC. Phosphorus in the culture of the Amazon River prawn (*Macrobrachium amazonicum*) and tambaqui (*Colossoma macropomum*) farmed in monoculture and in integrated multi-trophic systems. JWAS. 2019;1–22.

78. D’Alessandro ME, Collins PA. Manipulación nutricional en el camarón *Macrobrachium borellii* del río Paraná (Argentina) como recurso para la alimentación humana. Limnética. 2020; 39: 499–510.

79. Calvo NS, Reynoso CM, Resnik S, Cortés-Jacinto E, Collins P. Thermal stability of astaxanthin in oils for its use in fish food technology. Anim Feed Sci Tech. 2020; 270: 114668.

80. Chopin T, Robinson SMC, Troell M, Neori A, Buschmann AH, Fang J. Multitrophic integration for sustainable marine aquaculture. In: Ecological Engineering. Vol. 3 of Encyclopedia of Ecology. (Eds: S.E. Jørgensen & D.F. Brian). Oxford: Elsevier. 2008.

81. Vanni MJ, Layne CD. Nutrient recycling and herbivory as mechanisms in the ‘‘top-down’’ effect of fish on algae in lakes. Ecology. 1997;78:21–40.

82. Melo GAS. Manual de identificação dos crustáceos decápodos de água doce do Brasil. São Paulo: Editora Loyola. 2002.

83. Morrone JJ, Lopretto EC. Parsimony analysis of endemicity of freshwater Decapoda (Crustacea: Maclacostraca) from southern South America. Neotropica. 1995; 41: 3–8.

84. Torres MV, Giri F, Collins PA. Geometric morphometric analysis of the freshwater prawn *Macrobrachium borellii* (Decapoda: Palaemonidae) at a microgeographical scale in a floodplain system. Ecol Res. 2014; 29: 959–968.

85. Bond-Buckup G, Buckup L. A família Aeglidae (Crustacea, Decapoda, Anomura). Arq Zool. 1994; 32(4): 159–346.

86. Viau VE, López Greco LS, Bond-Buckup G, Rodríguez EM. Size at the onset of sexual maturity in the anomuran crab, *Aegla uruguayana* (Aeglidae). Acta Zool. 2006; 87(4): 253–264.

87. Diawol VP, Musin GE, Collins PA, Giri F. Distribution pattern in Aegla uruguayana Schmitt, 1942 (Decapoda: Anomura: Aeglidae): temporal and spatial approach. J Crust Biol. 2021; 41: 3 https://doi.org/10.1093/jcbiol/ruab038

88. Magalhães C. Famílias Pseudothelpulsidae e Trichodactylidae. In: Manual de identificação dos crustáceos decápodos de água doce do Brasil. (Ed: G. A. S. Melo) Loyola, São Paulo.

89. Torres MV, Giri, F, Collins PA. Temporal and spatial patterns of freshwater decapods associated with aquatic vegetation from floodplain rivers. Hydrobiologia. 2018; 823: 169–189.

90. Williner V, Collins PA. Feeding ecology of freshwater crab *Trichodactylus borellianus* (Decapoda: Trichodactylidae) in the floodplain of the Paraná River, southern South America. Lat Am J Aquat Res. 2013; 41: 781–792. DOI 10.3856/vol41-issue4-fulltext-15.

91. Carvalho DA, Collins PA, De Bonis CJ. The mark-recapture method applied to population estimates of a freshwater crab on an alluvial plain. Mar Freshw Res. 2013; 64: 317–323.

92. Devine JA, Vanni MJ. Spatial and seasonal variation in nutrient excretion by benthic invertebrates in a eutrophic reservoir. Freshw Biol. 2002; 47: 1107–1121.

93. National Research Council (NRC). Guide for the care and use of laboratory animals. The National Academy Press, Washington, D. C. 1996.

94. Koroleff F. Direct determination of ammonia in natural waters as indophenol blue. Information on techniques and methods for seawater analysis. An Interlab. report nr 3. Cons. Int. pour l’Exploration de la Mer. 1970; 3: 19 – 22.

95. Murphy J, Riley IP. A modified single solution method for the determination of phosphate in natural waters. Anal Chim Acta. 1962; 27: 31–36.

96. Andersen JM. An ignition method for determination of total phosphorous in lake sediments. Water Res. 1976; 10: 329–331.

97. R Development Core Team (2020). R: A language and environment for statistical computing. R Foundation for Statistical Computing, Vienna, Austria.

98. Kuznetsova, A., Brockhoff, P. B., & Christensen, R. H. B. (2015). Package ‘lmertest’. R package version, 2(0).

99. Lenth RV, Buerkner P, Herve M, Love J, Riebl H, Singmann H. Emmeans package: Estimated Marginal Means, aka Least-Squares Means. R package version >= 3.5.0 1.5.4. 2021.

100. Tanaka Y. Influence of body weight and food density on kinetics of ammonia excretion by the brine shrimp *Artemia franciscana*. JWAS. 1993; 24(4): 499–503.

101. Gido KB. Interspecific comparisons and the potential importance of nutrient excretion by benthic fishes in a large reservoir. Trans Am Fish Soc. 2002; 131: 260–270.

102. Schmidt-Nielsen K. Fisiologia animal – Adaptacão e meio ambiente. Cap.9. São Paulo: Santos. 2010.

103. Halvorson HM, Small GE. Observational field studies are not appropriate tests of consumer stoichiometric homeostasis. Freshw Sci, 2016; 35(4): 1103–1116.

104. El-Sabaawi RW, Warbanski ML, Rudman SM, Hovel R, Matthews B. Investment in boney defensive traits alters organismal stoichiometry and excretion in fish. Oecologia. 2016;181: 1209–1220. https://doi.org/10.1007/s00442-016-3599-0

105. Halvorson HM, Atkinson CL. Egestion versus excretion: a meta-analysis examining nutrient release rates and ratios across freshwater fauna. Diversity. 2018;11 (10), 189. https://doi.org/10.3390/d11100189

106. Viozzi MF, Martínez del Rio C, Williner V. Tissue-specific isotopic incorporation turnover rates and trophic discrimination factors in the freshwater shrimp Macrobrachium borellii (Crustacea: Decapoda: Palaemonidae). Zool Stud. 2021; 60: 32. doi:10.6620/ZS.2021.60-32

107. Kornberg A. Inorganic polyphosphate: toward making a forgotten polymer unforgettable. J Bacteriol. 1995; 177(3): 491–496.

108. Raubenheimer D, Jones SA. Nutritional imbalance in an extreme generalist omnivore: tolerance and recovery through complementary food selection. Anim Behav. 2006; 71: 1253–1262, doi:10.1016/j.anbehav.2005.07.024

109. Viozzi MF, Cabrera JM, Giri F, Carvalho DA, Williner V. Ontogenetic shifts in natural diet, chelae, and mandible of an omnivorous freshwater crab (*Aegla uruguayana*): linking morphology and function. Can J Zool. 2021; 99(8): 625–641.

110. Carvalho DA, Collins PA, Lima-Gomes R, Magalhães C, Torres MV, Williner V. A comparative study of the gastric ossicles of Trichodactylidae crabs (Brachyura: Decapoda) with comments on the role of diet and phylogeny in shaping morphological traits. PeerJ. 2018; 6:e5028; DOI 10.7717/peerj.5028.

111. Hendrixson HA, Sterner RW, Kay AD. Elemental stoichiometry of freshwater fishes in relation to phylogeny, allometry and ecology. J Fish Biol. 2007; 70: 121–140.

112. Vincent JFV. Arthropod cuticle: a natural composite shell system. Compos-A: Appl Sci Manuf. 2002; 33: 1311–1315.

113. Raabe D, Sachs C, Romano P. The crustacean exoskeleton as an example of a structurally and mechanically graded biological nanocomposite material. Acta Mater. 2005;53: 4281–4292.

114. Dillaman RM, Roer R, Shafer T, Modla S. The Crustacean integument: structure and function. In: Functional Morphology & Diversity. Vol. 1 (Eds: L. Watling & M. Thiel) Oxford University Press. 2013.

115. FAO. The State of World Fisheries and Aquaculture 2020. Sustainability in action. Roma. 2020. Available: https://www.fao.org/publications/card/es/c/CA9229EN

116. Holmer M, Black K, Duarte CM, Marbà N, Karakassis I. (Eds.). Aquaculture in the Ecosystem. Springer, Dordrecht, The Netherlands. 2008. 10.1017/CBO9781107415324.004

117. Chopin T. Integrated Multi-Trophic Aquaculture. Ancient, adaptable concept focuses on ecological integration. GAA. 2013; 16(2): 16–19.

118. Chopin T. Integrated Multi-Trophic Aquaculture (IMTA) is a concept, not a formula. International Aquafeed. February 2021, 18–19.

119. Shearer KD, Åsgård T, Andorsdöttir G, Aas GH. Whole body elemental and proximate composition of Atlantic salmon (*Salmo salar*) during the life cycle. J Fish Biol. 1994; 44 (5): 785–797.

120. Kaushik S, Georga I, Koumoundouros G. Growth and body composition of zebrafish (*Danio rerio*) larvae fed a compound feed from first feeding onward: toward implications on nutrient requirements. Zebrafish. 2011; 8 (2): 87–95.

